# Pharmacological CARD9 Inhibition Reduces *Pneumocystis murina* Burden In Vivo and Ex Vivo

**DOI:** 10.64898/2026.11.19.767846

**Authors:** Jenna L. McLean, Theodore J. Kottom, Andrew H. Limper

## Abstract

Caspase recruitment domain-containing protein 9 (CARD9) is an important adaptor protein of the innate immune responses to various fungal pathogens, including *Pneumocystis* species. CARD9 signaling is paramount in Pneumocystis pneumonia (PCP) host defense, and excessive inflammatory responses driven by this molecule can promote deleterious lung injury. We previously demonstrated that BRD5529, a first generation small-molecule CARD9 inhibitor, attenuates pulmonary inflammatory cytokine production during PCP. Here, we investigated the previously unreported effects of BRD5529 on *Pneumocystis murina* burden in vivo and in an ex vivo PCP-infected precision-cut lung slice (PCLS) model. In vitro, BRD5529 treatment (1 mg/kg) reduced lung organism burden in mice with established PCP, with an effect similar in magnitude to standard-of-care trimethoprim-sulfamethoxazole (TMP-SMX) therapy. In infected PCLS cultured with BRD5529 for 72 h, *P. murina* viability was also reduced, with the greatest reduction observed at lower BRD5529 concentrations. BRD5529 treatment did not reduce PCLS metabolic activity and viability after 72 h, as assessed by resazurin reduction. Together, these findings suggest that pharmacological CARD9 inhibition may impact organism burden/viability in addition to reducing lung inflammation during PCP.

**IMPORTANCE:** Pneumocystis pneumonia (PCP) remains a serious opportunistic infection in immunocompromised individuals. Although inflammatory responses aid in organism clearance, excessive inflammation can promote lung injury. We have previously shown that the CARD9 inhibitor BRD5529 reduces pulmonary inflammation in a CD4 depleted mouse model of PCP. Here, we report the unexpected observation that BRD5529 also reduces *P. murina* burden in infected mice and decreases fungal viability in infected lung tissue-maintained ex vivo. These findings may suggest that pharmacological targeting of CARD9-assocated signaling may influence both lung inflammatory responses and organism control during PCP.

## OBSERVATION

*Pneumocystis jirovecii* pneumonia (PJP) remains an important cause of morbidity and mortality among immunocompromised individuals (1-3). Although TMP-SMX remains the standard preferred treatment, drug-associated toxicity and treatment intolerance highlight the need for alternative therapeutic approaches (4). During PCP, lung injury is associated with exuberant host inflammatory response to the organism (5). Innate immune recognition of *Pneumocystis* involves several pattern-recognition receptors, including C-type lectin receptors such as Dectin-1/Dectin-2, which can signal through the adapter protein CARD9 (6, 7). In particular, β-glucans present in the *Pneumocystis* cell wall activate Dectin-1-associated signaling pathways, promoting downstream inflammatory responses (8-10). Although CARD9-dependent signaling is important for macrophage activation and antifungal host defense, excessive activation of these pathways may contribute to pulmonary inflammation and tissue injury during PCP (11). BRD5529 is a first-generation small-molecule inhibitor of CARD9-associated inflammatory signaling (12). Previous studies demonstrated that BRD5529 attenuates macrophage inflammatory responses to *Pneumocystis* β-glucans (13, 14). More recently, we demonstrated that BRD5529 significantly reduces IL-6, TNF-α, and IL-1β during PCP (15). However, the effects of BRD5529 on lung *Pneumocystis* burden have not been previously described. Accordingly, we investigated whether BRD5529 treatment influences *P. murina* burden during PCP and whether its effects could be reproduced in an ex vivo PCP PCLS model. We previously demonstrated that BRD5529 significantly attenuates lung inflammatory cytokine production in mice with established PCP (15). In those experiments, CD4-depleted mice infected with *P. murina* for seven weeks received BRD5529 (1mg/kg) or vehicle by daily intraperitoneal injection for 10 days. To determine whether CARD9 inhibition also influenced organism burden, we evaluated lung *P. murina* burdens in these animals. An additional cohort of mice with established PCP received therapeutic TMP-SMX (200 mg/L trimethoprim and 1,000 mg/L sulfamethoxazole (16) as a standard of care treatment during the same treatment period.

BRD5529 treatment significantly reduced lung fungal burden compared with infected controls (Fig. 1). The magnitude of this reduction was similar to that observed in mice receiving therapeutic TMP-SMX. These findings were unexpected because CARD9 contributes to innate antifungal host defense, and genetic CARD9 deficiency has previously been associated with impaired *Pneumocystis* clearance. Our observations suggest that pharmacological inhibition of CARD9-associated inflammatory signaling during established infection may have effects on organism burden that differ from those associated with complete genetic loss of CARD9 infection. Because BRD5529 reduced lung organism burden in vivo, we next investigated whether this effect could be reproduced in infected lung tissue-maintained ex vivo. *P. murina*-infected PCLS (5 mm in diameter and 300 um thick) were exposed to BRD5529 (1, 10, or 50 μM), pentamidine (1 μM), or vehicle for 72 h. Organism viability was determined by reverse transcription-quantitative PCR targeting *P. murina mitochondrial16S* rRNA. BRD5529 reduced organism viability in infected PCLS after 72 h, with the greatest reduction observed at 1 μM (Fig. 2A). Interestingly, increasing BRD5529 concentrations did not result in progressively greater reductions in *P. murina mitochondrial 16S* rRNA signal. To determine whether these effects were associated with generalized tissue toxicity, PCLS metabolic activity was assessed by resazurin following 72 h of BRD5529 exposure. BRD5529 did not reduce PCLS viability across the concentrations tested (Fig. 2B). Expression of the type I and type II alveolar epithelial cell-associated genes *Pdpn* and *Sftpc*, respectively, was also assessed after 72 h to characterize epithelial responses to treatment and potential changes in alveolar epithelial phenotype (Fig. S1). Together with preserved PCLS metabolic activity, these findings suggest that the BRD5529-associated reduction in *P. murina* viability was not simply attributable to overt PCLS toxicity. The observation that BRD5529 reduces *P. murina* burden while suppressing lung inflammatory responses raises important questions regarding the relationship between CARD9 signaling, inflammation, and organism clearance. Although CARD9-dependent signaling contributes to antifungal host defense, excessive inflammatory responses during PCP also promote lung injury. These findings suggest that pharmacological modulation of inflammatory signaling may have consequences for organism control that differ from those associated with complete disruption of protective immune pathways. Previous studies have demonstrated that the host inflammatory response to *Pneumocystis* plays a dual role during infection, contributing to both organism clearance and lung injury (17, 18). Chen et al. demonstrated that TNF-α is important for effective *Pneumocystis* clearance (19), whereas Wright et al. showed that excessive immune-mediated inflammation contributes substantially to pulmonary dysfunction (17). Pharmacological immunomodulation provides additional evidence that these processes can, under some conditions, be disassociated. Treatment with sulfasalazine has been shown to attenuate *Pneumocystis*-associated pulmonary inflammation while enhancing macrophage-mediated organism clearance. Collectively, these findings highlight the importance of maintaining a balance between protective immunity and excessive inflammation during *Pneumocystis* infection. They further suggest that selective attenuation of inflammatory signaling can, under some conditions, improve organism control, whereas disruption of pathways required for protective immunity can compromise organism clearance (18, 20).

**FIG 1.**
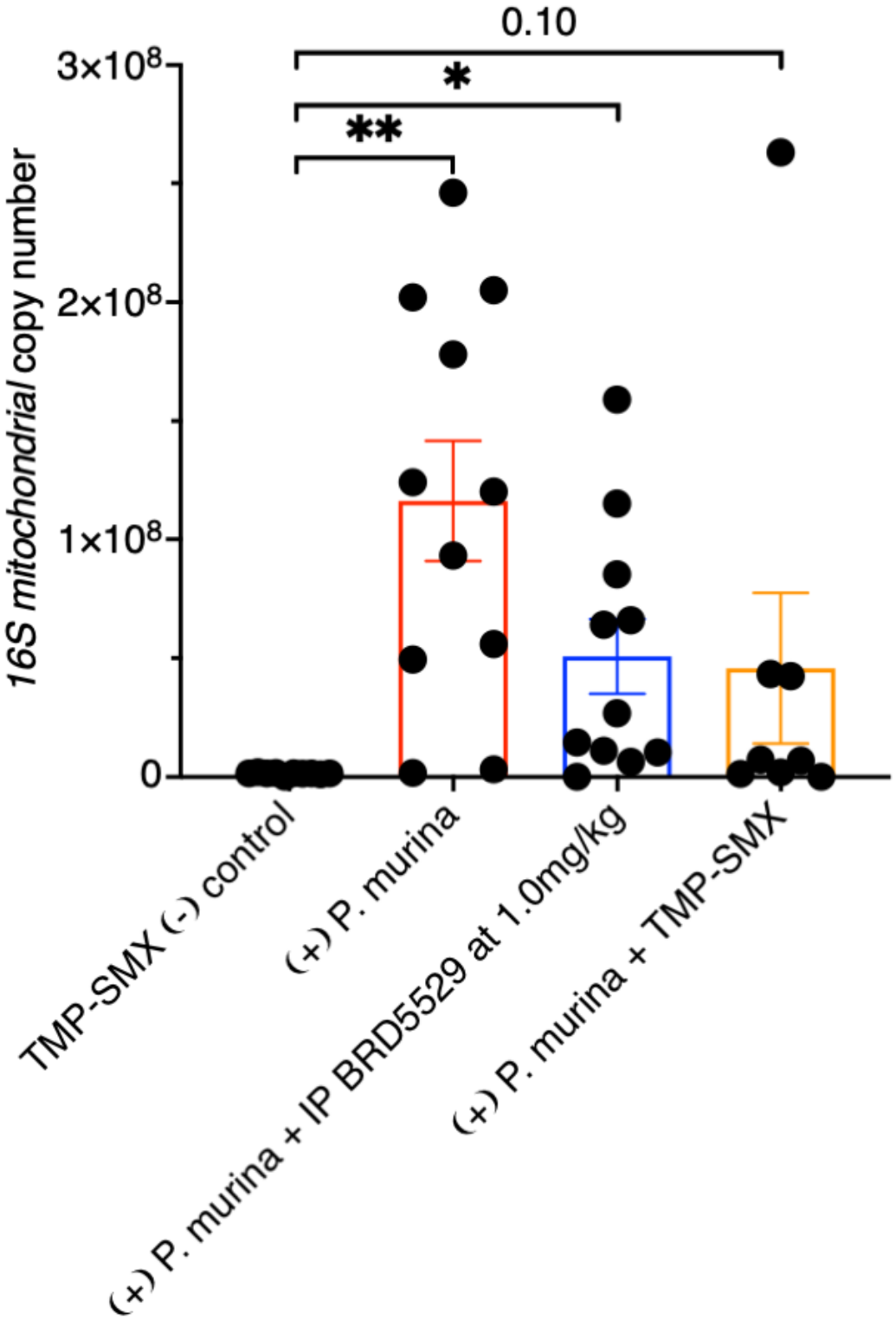
BRD5529 reduces lung *P. murina* burden during established PCP. Balb/c Mice were depleted of CD4-positive lymphocytes with GK1.5 and infected with *P. murina* (∼1 x 10^6^ organisms/mouse) as previously described (15). Following 7 weeks of established infection, mice received BRD5529 (1mg/kg), formulated in 1% Methocel, or vehicle by daily intraperitoneal injections for 10 days. A separate cohort of infected mice received therapeutic trimethoprim-sulfamethoxazole (TMP-SMX; 200 mg/L trimethoprim and 1000 mg/L sulfamethoxazole) in drinking water during the same 10-day treatment period using the regimen described by Powles et al. (16). Following treatment, lungs were collected, and *P. murina* burden was quantified by real-time qPCR targeting the *mitochondrial 16S* rRNA gene and determining copy number using previously established methods (23, 24). Individual symbols represent individual mice, and data are presented +/-SEM. Differences among treatment groups were analyzed by one-way ANOVA followed by multiple-comparisons test using GraphPad version 11.0.2 (GraphPad Software, San Diego, CA). *P* < 0.05 was considered statistically significant. Lung inflammatory cytokine measurements from the BRD5529-treated animals were reported previously (15); the organism-burden measurements presented here have not been previously published.

**FIG 2.**
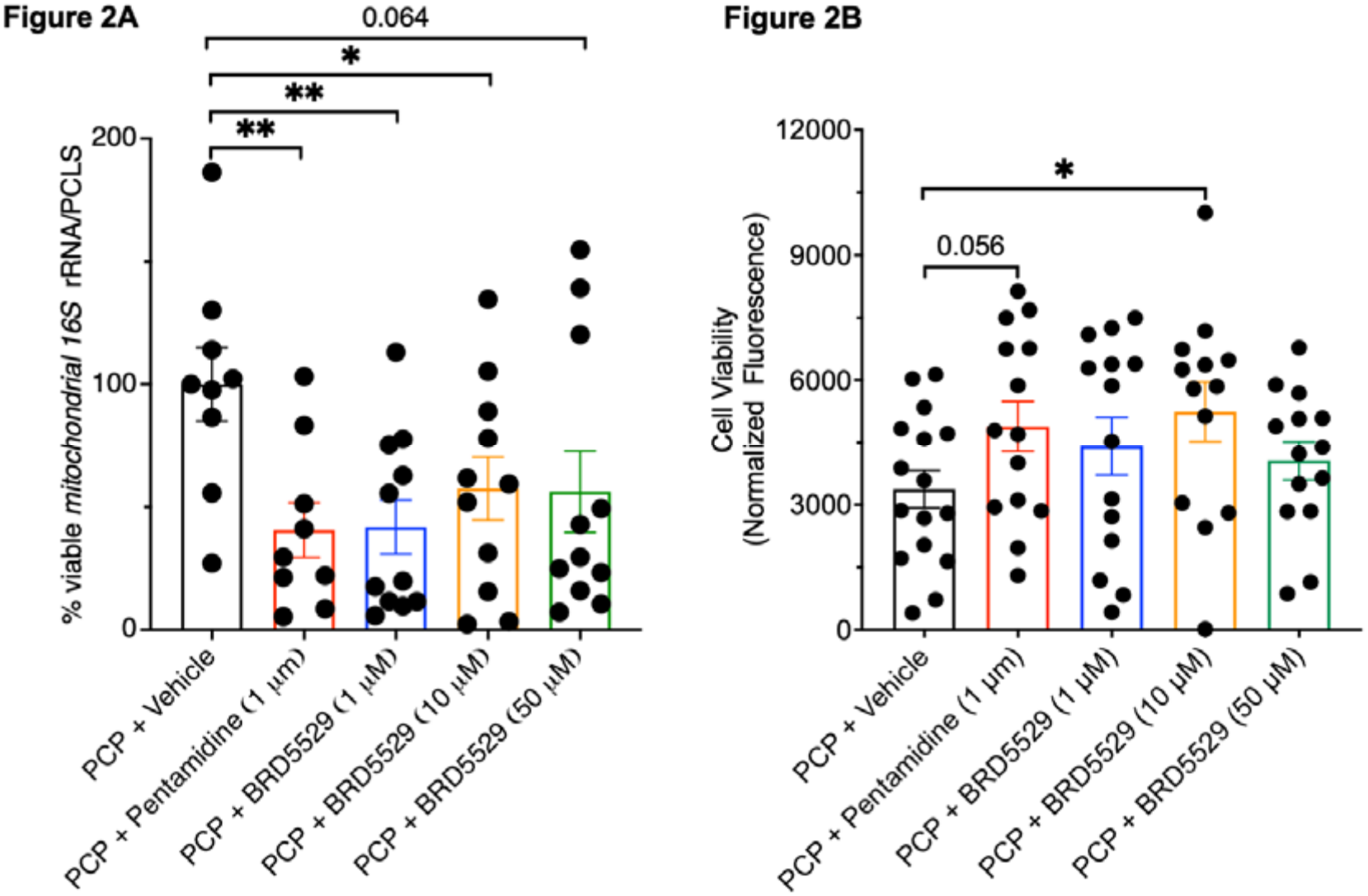
Effects of BRD5529 on *P. murina* viability-associated *mitochondrial 16S* rRNA and PCLS metabolic activity. *P. murina*-infected precision-cut lung slices (PCLS) were prepared and maintained using previously established methods (25) and cultured for 72 h with DMSO vehicle, pentamidine (1 μM), and BRD5529 (1, 10, or 50 μM). BRD5529 concentrations were selected to span the previously reported biochemical IC_50_ of 8.6 μM for inhibition of the CARD9-TRIM62 interaction (12). This concentration range also encompasses the approximately 10.7 μM theoretical concentration previously used to support selection of the 1mg/kg BRD5529 dose in murine studies (15). **(A)** Following treatment, total RNA was isolated from PCLS and reverse-transcribed to cDNA. *P. murina* viability was quantified by *mitochondrial 16S* rRNA qPCR using a standard curve generated from a purified 165-bp *mitochondrial 16S* amplicon. Values from individual PCLS were normalized to the mean *mitochondrial 16S* rRNA signal from the corresponding mouse-matched DMSO-treated PCP PCLS, which was defined as 100%. Pentamidine was included as an anti-*Pneumocystis* positive control. **(B)** PCLS metabolic activity was assessed after 72 h of treatment using a resazurin-based fluorescence assay and reported as fluorescence intensity (25). Individual points represent measurements from individual PCLS, with multiple slices obtained from each infected mouse. A total of 4-5 PCP infected mice were used. Data are presented as mean +/-SEM. Statistical analyses were performed using GraphPad Prism 11.0.2. and *P* < 0.05 was considered statistically significant.

This balance may also be particularly relevant to CARD9 signaling. We previously demonstrated that genetic CARD9 deficiency impairs host defense against *Pneumocystis*, indicating that CARD9-dependent responses contribute to effective organism clearance (11). In contrast, the present findings demonstrate that pharmacological CARD9 inhibition with BRD5529 during established infection is associated with reduced *P. murina* burden. Complete genetic absence of CARD9 throughout infection and therapeutic inhibition during established PCP represents fundamentally different perturbations of this pathway. Differences in the timing, magnitude, and cellular distribution of CARD9 inhibition could therefore influence the balance between protective antifungal responses and inflammatory lung injury. In this context, the differential response observed across BRD5529 concentrations in infected PCLS is intriguing. The greatest reduction in *P. murina* viability occurred at the lowest BRD5529 concentration examined, whereas higher concentrations produced less pronounced effects. One possible explanation is that BRD5529 differentially influences inflammatory signaling and macrophage-mediated antifungal infections. Previous studies have shown that pharmacological inhibition of inflammatory signaling during PCP can enhance macrophage-mediated organism clearance, suggesting that inflammatory responses and protective antifungal functions can, under some conditions, be modulated independently (15, 21). Furthermore, we previously demonstrated that BRD5529 inhibits *Pneumocystis* β-glucan-induced macrophage MAPK activation and TNF-α production in a concentration-dependent manner (14). Together, these observations raise the possibility that lower BRD5529 concentrations attenuate inflammatory signaling while preserving macrophage functions involved in organism clearance, whereas higher concentrations may have broader effects on protective host responses. However, macrophage function and CARD9 pathway activity were not directly evaluated across BRD5529 concentrations in the present study. Therefore, understanding the differential response to BRD5529 remains to be fully determined.

## ETHICAL STATEMENT

This study was performed in strict accordance with the recommendations in the Guide for the Care and Use of Laboratory Animals (22). The Mayo Clinic Institutional Animal Care and Use Committee approved this research (A00008661-26).

## FIGURE LEGENDS

**FIG S1.**
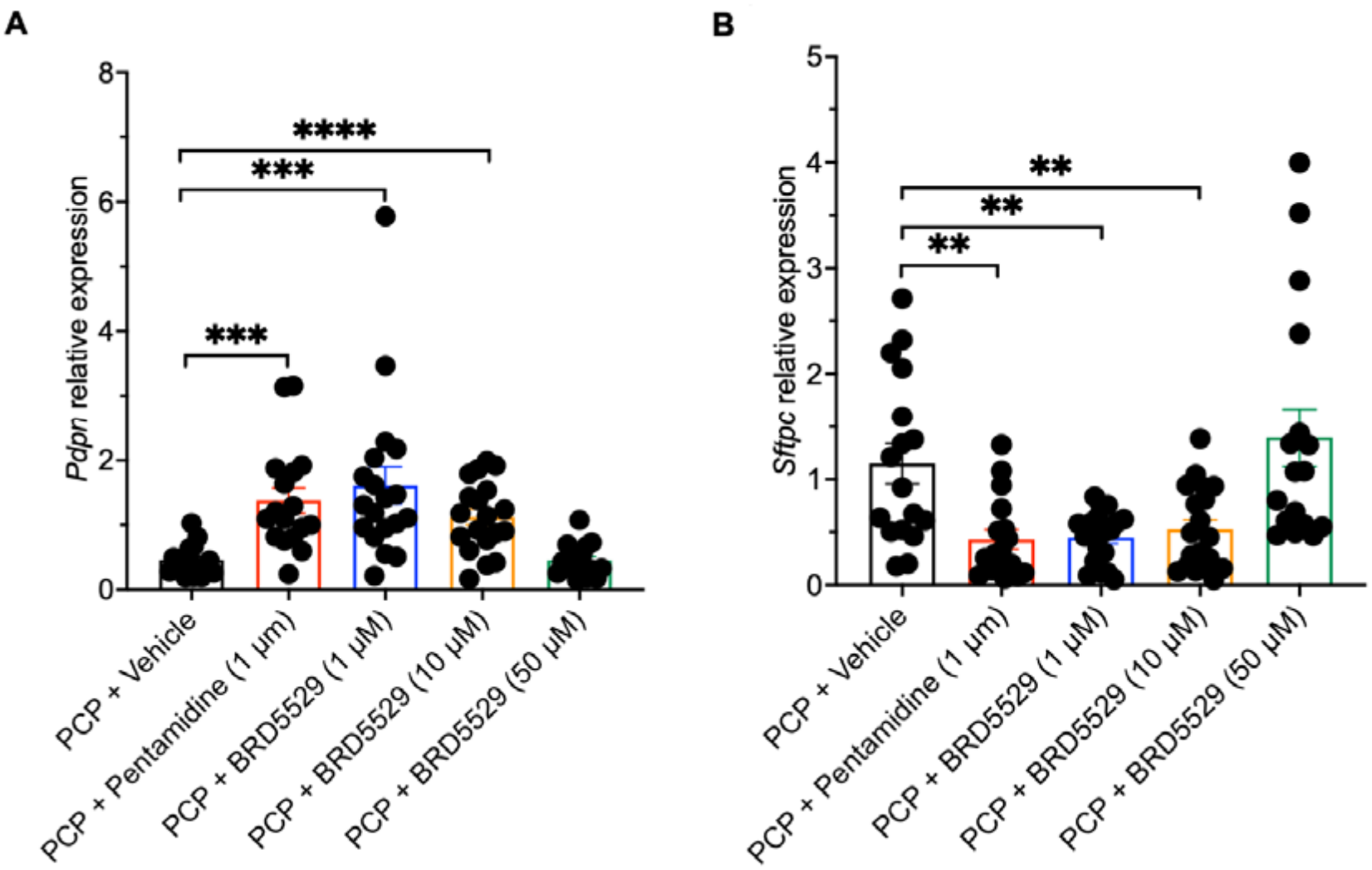
Effects of BRD5529 on epithelial-associated gene expression in *P. murina*-infected PCLS. *P. murina*-infected PCLS were cultured for 72 h with DMSO vehicle, pentamidine (1 μM), or BRD5529 (1, 10, 50 μM). Relative expression of **(A)** *Pdpn* and **(B)** *Sftpc* was determined using the 2^-ΔΔCt method, with *B2m* as the reference gene (26). Individual points represent measurements from individual PCLS obtained from five infected mice, with multiple slices analyzed per mouse. Mice were considered independent biological replicates. Data are presented as mean +/-SEM. Statistical analyses were conducted as previously noted. *P* < 0.05 was considered significant.

## REFERENCES

1. Haensel L, Sprute R, Grothe J, Robert-Gangneux F, Gangneux JP, Meis JF, Cornely OA, Koehler P. 2025. The 2025 EQUAL Pneumocystis Score-an ECMM tool to measure QUALity in Pneumocystis pneumonia management. JAC Antimicrob Resist 7:dlaf165.

2. Kamel T, Janssen-Langenstein R, Quelven Q, Chelly J, Valette X, Le MP, Bourenne J, Garot D, Fillatre P, Labruyere M, Heming N, Lambiotte F, Lascarrou JB, Lesieur O, Bachoumas K, Ferre A, Maury E, Chalumeau-Lemoine L, Bougon D, Roux D, Guisset O, Coudroy R, Boulain T, group P-MS. 2024. Pneumocystis pneumonia in intensive care: clinical spectrum, prophylaxis patterns, antibiotic treatment delay impact, and role of corticosteroids. A French multicentre prospective cohort study. Intensive Care Med 50:1228–1239.

3. Li J, Mu X, Li H, Liu X. 2025. Immune responses of different hosts to Pneumocystis infection. Eur Respir Rev 34.

4. McDonald EG, Butler-Laporte G, Del Corpo O, Hsu JM, Lawandi A, Senecal J, Sohani ZN, Cheng MP, Lee TC. 2021. On the Treatment of Pneumocystis jirovecii Pneumonia: Current Practice Based on Outdated Evidence. Open Forum Infect Dis 8:ofab545.

5. Kottom TJ, Limper AH. 2026. Modulation of host lung immunity in Pneumocystis pneumonia: a review of current concepts and future prospects for novel adjunctive immune-based therapies. J Immunol 215.

6. Kottom TJ, Carmona EM, Limper AH. 2021. Current State of Carbohydrate Recognition and C-Type Lectin Receptors in Pneumocystis Innate Immunity. Front Immunol 12:798214.

7. Kottom TJ, Hebrink DM, Jenson PE, Marsolek PL, Wuthrich M, Wang H, Klein B, Yamasaki S, Limper AH. 2018. Dectin-2 Is a C-Type Lectin Receptor that Recognizes Pneumocystis and Participates in Innate Immune Responses. Am J Respir Cell Mol Biol 58:232–240.

8. Ricks DM, Chen K, Zheng M, Steele C, Kolls JK. 2013. Dectin immunoadhesins and pneumocystis pneumonia. Infect Immun 81:3451–62.

9. Saijo S, Fujikado N, Furuta T, Chung SH, Kotaki H, Seki K, Sudo K, Akira S, Adachi Y, Ohno N, Kinjo T, Nakamura K, Kawakami K, Iwakura Y. 2007. Dectin-1 is required for host defense against Pneumocystis carinii but not against Candida albicans. Nat Immunol 8:39–46.

10. Steele C, Marrero L, Swain S, Harmsen AG, Zheng M, Brown GD, Gordon S, Shellito JE, Kolls JK. 2003. Alveolar macrophage-mediated killing of Pneumocystis carinii f. sp.muris involves molecular recognition by the Dectin-1 beta-glucan receptor. J Exp Med 198:1677–88.

11. Kottom TJ, Nandakumar V, Hebrink DM, Carmona EM, Limper AH. 2020. A critical role for CARD9 in pneumocystis pneumonia host defence. Cell Microbiol 22:e13235.

12. Leshchiner ES, Rush JS, Durney MA, Cao Z, Dancik V, Chittick B, Wu H, Petrone A, Bittker JA, Phillips A, Perez JR, Shamji AF, Kaushik VK, Daly MJ, Graham DB, Schreiber SL, Xavier RJ. 2017. Small-molecule inhibitors directly target CARD9 and mimic its protective variant in inflammatory bowel disease. Proc Natl Acad Sci U S A 114:11392–11397.

13. Kottom TJ, Schaefbauer K, Carmona EM, Yi ES, Limper AH. 2022. Preclinical and Toxicology Studies of BRD5529, a Selective Inhibitor of CARD9. Drugs R D 22:165–173.

14. Kottom TJ, Carmona EM, Limper AH. 2020. Targeting CARD9 with Small-Molecule Therapeutics Inhibits Innate Immune Signaling and Inflammatory Response to Pneumocystis carinii beta-Glucans. Antimicrob Agents Chemother 64.

15. Kottom TJ, Carmona EM, Schaefbauer K, Stelzig KE, Pellegrino MR, Bindzus M, Limper AH. 2025. The importance of Fcgamma and C-type lectin receptors in host immune responses during Pneumocystis pneumonia. Infect Immun 93:e0027624.

16. Powles MA, Liberator P, Anderson J, Karkhanis Y, Dropinski JF, Bouffard FA, Balkovec JM, Fujioka H, Aikawa M, McFadden D, Schmatz D. 1998. Efficacy of MK-991 (L-743,872), a semisynthetic pneumocandin, in murine models of Pneumocystis carinii. Antimicrob Agents Chemother 42:1985–9.

17. Bhagwat SP, Gigliotti F, Xu H, Wright TW. 2006. Contribution of T cell subsets to the pathophysiology of Pneumocystis-related immunorestitution disease. Am J Physiol Lung Cell Mol Physiol 291:L1256–66.

18. Zhang ZQ, Gigliotti F, Wright TW. 2023. The Dual Benefit of Sulfasalazine on Pneumocystis Pneumonia-Related Immunopathogenesis and Antifungal Host Defense Does Not Require IL-4Ralpha-Dependent Macrophage Polarization. Infect Immun 91:e0049022.

19. Chen W, Havell EA, Harmsen AG. 1992. Importance of endogenous tumor necrosis factor alpha and gamma interferon in host resistance against Pneumocystis carinii infection. Infect Immun 60:1279–84.

20. Wang J, Gigliotti F, Bhagwat SP, George TC, Wright TW. 2010. Immune modulation with sulfasalazine attenuates immunopathogenesis but enhances macrophage-mediated fungal clearance during Pneumocystis pneumonia. PLoS Pathog 6:e1001058.

21. Zhang ZQ, Wang J, Hoy Z, Keegan A, Bhagwat S, Gigliotti F, Wright TW. 2015. Neither classical nor alternative macrophage activation is required for Pneumocystis clearance during immune reconstitution inflammatory syndrome. Infect Immun 83:4594–603.

22. (ed). 2011. Guide for the care and use of laboratory animals. National Academies Press,, Washington, D.C. http://www.ncbi.nlm.nih.gov/books/NBK54050ebrary http://site.ebrary.com/id/10443276 National Academies Press http://www.nap.edu/catalog.php?record_id=12910 National Academies Press http://www.nap.edu/catalog.php?record_id=12910#toc http://www.ncbi.nlm.nih.gov/bookshelf/br.fcgi?book=nap12910 http://grants.nih.gov/grants/olaw/Guide-for-the-Care-and-use-of-laboratory-animals.pdf. Accessed

23. Wakefield AE, Pixley FJ, Banerji S, Sinclair K, Miller RF, Moxon ER, Hopkin JM. 1990. Detection of Pneumocystis carinii with DNA amplification. Lancet 336:451–3.

24. Kottom TJ, Hebrink DM, Jenson PE, Nandakumar V, Wuthrich M, Wang H, Klein B, Yamasaki S, Lepenies B, Limper AH. 2017. The Interaction of Pneumocystis with the C-Type Lectin Receptor Mincle Exerts a Significant Role in Host Defense against Infection. J Immunol 198:3515–3525.

25. Pellegrino M, Kottom TJ, Achilonu CC, Stelzig KE, Ayyalasomayajula S, Gibbard DF, Cai J, Brownfield DG, Limper AH. Accepted for publication. Ex Vivo Modeling of Pneumocystis murina Infection Using Precision-Cut Lung Slices to Evaluate Drug Susceptibility and Host Immune Responses. The Journal of Infectious Diseases.

26. Livak KJ, Schmittgen TD. 2001. Analysis of relative gene expression data using real-time quantitative PCR and the 2(-Delta Delta C(T)) Method. Methods 25:402–8.

